# Fractional-Order Temporal Dynamics of Tumour–Immune–Chemotherapy Interactions: A Reproducible Dimensionless Analysis and a Pharmacokinetic Caveat

**DOI:** 10.64898/2026.09.29.755373

**Authors:** Fatemeh Yadollahzadeh Chari, Amin Gholami Davodi

**Affiliations:** Babol University of Medical Sciences; Babol Noshirvani University of Technology

**Keywords:** fractional differential equations, Caputo derivative, tumour–immune dynamics, pharmacokinetics, Mittag–Leffler relaxation, predictor–corrector method, reproducibility

## Abstract

Fractional differential equations are increasingly used to represent temporal memory in tumour–immune and pharmacokinetic models, but changing the derivative order without a consistent time scale can make comparisons difficult to interpret. We study a dimensionless, commensurate Caputo system for tumour, immune, normal-cell, and drug states. We establish positivity, boundedness, the tumour-invasion threshold, and local stability conditions. We then use a fractional Adams–Bashforth–Moulton predictor–corrector method with an explicit mesh-convergence study. Two conclusions correct potentially misleading interpretations of the model. First, the equilibrium locations are independent of the fractional order: decreasing the order changes transient time scales but does not by itself alter the model’s asymptotic states. Second, after a unit bolus the drug equation has the exact solution *C*(*τ*) = *E*_*α*_(*− kτ* ^*α*^). Its maximum is therefore the initial concentration, not a delayed peak. For 0 *< α <* 1, the algebraic tail makes the infinite-horizon area under the concentration curve diverge. In the illustrative coupled simulations, the apparent late tumour reduction at small *α* is consequently driven by persistent modelled drug exposure and must not be read as evidence of superior therapy. All parameters are dimensionless and illustrative; no patient data are fitted. Source code and generated tables are included to make every numerical statement reproducible.

## 1 Introduction

Mathematical models of tumour–immune interactions provide a controlled setting in which feedback, immune escape, and treatment schedules can be examined before more detailed experimental questions are posed. Classical systems of ordinary differential equations have been used to describe tumour growth, immune surveillance, and the combined effects of immunotherapy and chemotherapy [1, 2]. Such models are necessarily simplified, but they help separate mechanisms that may otherwise be difficult to reason about.

Ordinary differential equations are Markovian at the level of the chosen state: once the present state is known, its previous trajectory does not enter explicitly. Fractional derivatives provide one way to introduce nonlocal temporal dependence. In particular, the Caputo derivative weights the entire past through a power-law kernel [3, 4]. Fractional tumour–immune systems and associated optimal-control problems have therefore received considerable attention [5, 6]. Related fractional constructions have also been studied in pharmacokinetics and pharmacodynamics [7–9].

The formal replacement of a first derivative by a derivative of order *α* is not innocuous. A derivative with respect to dimensional time has units of time^−*α*^, so rate coefficients taken unchanged from an integer-order model no longer have the same units. Moreover, changes in a finite-time output can reflect a slower transient rather than a change in equilibrium. In a fractional clearance equation, a small value of *α* also produces a heavy concentration tail. If these points are not made explicit, a numerical comparison may be assigned a biological or therapeutic meaning that the equations do not support.

This paper gives a transparent analysis of a phenomenological four-state model inspired by established tumour–immune–drug models. The contribution is methodological rather than clinical. We (i) formulate the system in dimensionless time, (ii) derive the correct tumour-invasion threshold and stability conditions, (iii) show that equilibria do not depend on *α*, (iv) identify the exact Mittag–Leffler pharmacokinetic solution and its non-integrable tail for *α <* 1, and (v) provide reproducible computations with mesh refinement. We deliberately omit patient subgroup, age, sex, and circadian claims because the model is not calibrated to data that could support them. We likewise do not report likelihood-based model selection without an identified dataset and observation model.

## 2 Dimensionless fractional model

### 2.1 Caputo derivative and time scaling

For 0 *< α <* 1, the Caputo derivative of an absolutely continuous function *x* is

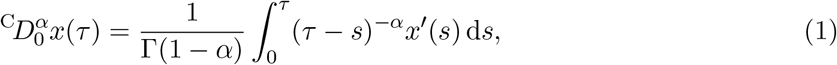

and 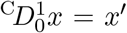. The dimensionless variable *τ* = *t/t*_ref_ is used throughout. Consequently, every coefficient below is dimensionless. A fitted physical-time model would require a declared reference time and an explicit map between dimensionless and dimensional coefficients. Without such a map, numerical values of *τ* must not be labelled as days or hours.

### 2.2 State equations

Let *T, I, N*, and *C* denote non-negative, dimensionless tumour, immune, normal-cell, and drug states. We study the commensurate system

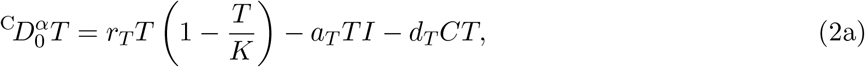

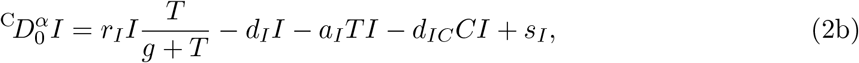

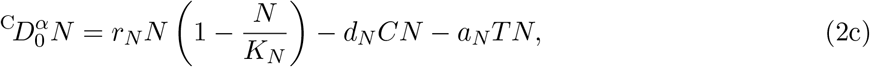

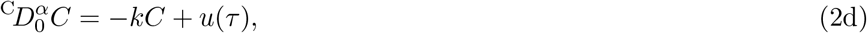

where 0 *< α* ≤ 1. Tumour and normal cells have logistic growth; immune cells receive a constant source and a saturating tumour-dependent stimulation; tumour and drug suppress immune cells; and the drug damages tumour and normal cells. The system is phenomenological. It is related in structure, but not identical, to the validated models in de Pillis et al. [1, 2]; our numerical coefficients must therefore be viewed as illustrative.

Table 1 lists the coefficients used in the simulations. Two regimes differ only in tumour killing by immune cells. The tumour-escape regime uses *a*_*T*_ = 0.10, whereas the immune-control regime uses *a*_*T*_ = 0.50. Initial conditions for untreated simulations are (*T*_0_, *I*_0_, *N*_0_, *C*_0_) = (0.05, 0.20, 1, 0). A unit bolus is represented by *C*_0_ = 1 and *u*(*τ*) = 0 for *τ >* 0; treating a bolus as the ordinary function *u*(0) = 1 would have no finite impulse mass.

**Table 1:** Dimensionless coefficients used in the illustrative simulations.

| Coefficient | Value | Interpretation |
| --- | --- | --- |
| $r_T$ | 0.15 | tumour growth |
| $K$ | 1.00 | tumour carrying level |
| $a_T$ | 0.10 or 0.50 | immune-mediated tumour killing |
| $d_T$ | 0.30 | drug effect on tumour |
| $r_I$ | 0.20 | immune stimulation |
| $g$ | 0.30 | immune half-saturation level |
| $d_I$ | 0.10 | immune loss |
| $a_I$ | 0.40 | tumour-mediated immune suppression |
| $d_{IC}$ | 0.20 | drug-mediated immune suppression |
| $s_I$ | 0.05 | basal immune source |
| $r_N$ | 0.10 | normal-cell growth |
| $K_N$ | 1.00 | normal-cell carrying level |
| $d_N$ | 0.25 | drug damage to normal cells |
| $a_N$ | 0.10 | tumour competition with normal cells |
| $k$ | 0.20 | drug clearance coefficient |

## 3 Analytical properties

### 3.1 Positivity and boundedness

#### Proposition 1

(Positive invariance). *Suppose the initial state is non-negative and u*(*τ*) ≥ 0. *Then every solution of* (2), *while it exists, remains in* 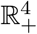.

*Proof*. The vector field is locally Lipschitz on the non-negative orthant. On the boundary faces *T* = 0, *I* = 0, *N* = 0, and *C* = 0, the corresponding right-hand sides are respectively 0, *s*_*I*_ *>* 0, 0, and *u*(*τ*) ≥ 0. Thus the vector field is quasipositive. The fractional comparison principle then gives positive invariance; see, for example, Diethelm [4, Chapters 6–7].

#### Proposition 2

(Uniform bounds). *For non-negative initial conditions and a bounded non-negative input, the states of* (2) *are bounded. In particular*,

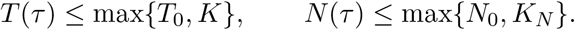

*If*

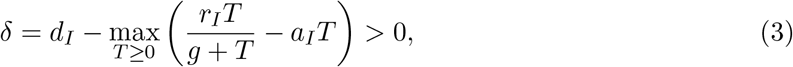

*then I*(*τ*) ≤ max*{I*_0_, *s*_*I*_*/δ}. The parameters in Table 1 give δ* ≈ 0.0898.

*Proof*. Dropping non-positive terms in (2a) and (2c) gives scalar logistic upper comparisons. The coefficient multiplying *I* in (2b) is no larger than −*δ* because *C* ≥ 0. Hence 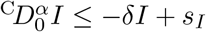. The stable linear drug equation is bounded for bounded input. Applying the scalar comparison principle yields the stated estimates.

The maximum in (3) is zero when *r*_*I*_ ≤ *a*_*I*_*g* and otherwise equals 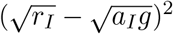. This gives an explicit, easily checked sufficient condition for a uniform immune bound.

### 3.2 Equilibria and tumour invasion

For *u* = 0, the tumour-free equilibrium is

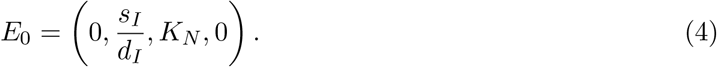

Because the Caputo derivative of a constant is zero, equilibrium locations are independent of *α*.

Define the tumour-invasion number

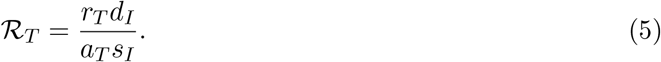

The tumour-direction eigenvalue at *E*_0_ is

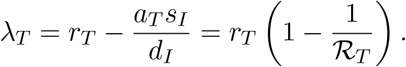

The remaining eigenvalues are −*d*_*I*_, −*r*_*N*_, and −*k*. For a commensurate Caputo system, local asymptotic stability requires | arg(*λ*)| *> απ/*2 for every eigenvalue [10]. All four eigenvalues are real, so *E*_0_ is locally asymptotically stable for every 0 *< α* ≤ 1 if *ℛ*_*T*_ *<* 1, and unstable if *ℛ*_*T*_ *>* 1. The illustrative regimes have *ℛ*_*T*_ = 3.0 and 0.6, respectively.

At a positive untreated equilibrium,

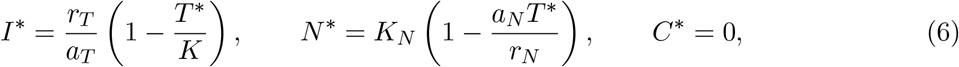

and *T* ^∗^ solves

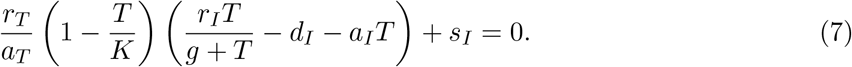

For the escape regime, the admissible solution is

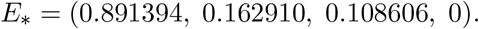

The Jacobian eigenvalues at this point are approximately −0.10764, −0.33299, −0.01086, and −0.20. It is therefore locally asymptotically stable for every tested fractional order, and indeed for every 0 *< α* ≤ 1. This calculation makes the distinction between equilibrium and transient effects explicit: varying *α* changes the route to *E*_∗_, not *E*_∗_ itself.

### 3.3 Exact fractional clearance and its consequence

For a unit bolus, *C*(0) = 1 and *u* = 0, equation (2d) has the exact solution

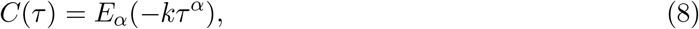

where 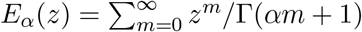 is the Mittag–Leffler function. For 0 *< α* ≤ 1 and *k >* 0, (8) is completely monotone. It follows immediately that

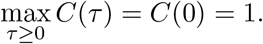

Thus this equation cannot produce a delayed concentration peak after an instantaneous bolus.

For 0 *< α <* 1, the large-time asymptotic form is

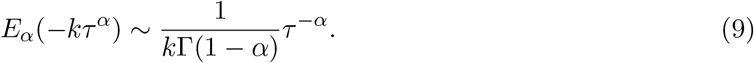

Consequently,

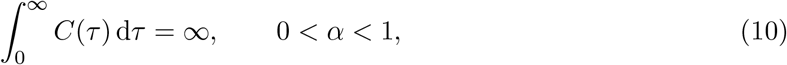

whereas the integer-order value is 1*/k*. A finite-window area under the curve is well defined, but it depends on the chosen horizon and should not be interpreted as conventional total exposure without qualification. Fractional pharmacokinetic models can be useful [7–9]; equation (10) simply shows that the one-equation construction used here has a strong structural implication.

## 4 Numerical method and reproducibility

We solve (2) with the fractional Adams–Bashforth–Moulton predictor–corrector method of Diethelm et al. [11]. Let *τ*_*n*_ = *nh*. For a vector equation 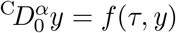, the predictor is

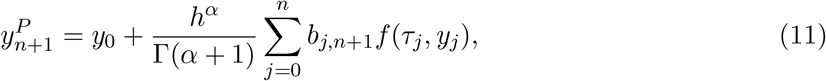

where *b*_*j,n*+1_ = (*n* + 1 − *j*)^*α*^ − (*n* − *j*)^*α*^. The corrector is

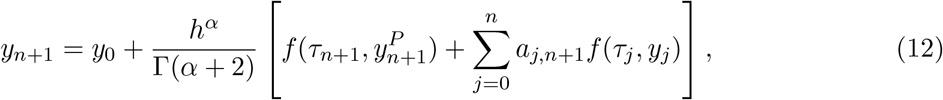

with the standard fractional Adams weights [11, 12]. Main experiments use *h* = 0.05. The code uses no curve smoothing and writes the plotted values and summary metrics directly from the solver.

Table 2 compares terminal states at *τ* = 20 with a reference computation using *h* = 0.00625. The empirical rate is close to 1 + *α* for *α* = 0.7 and to second order for *α* = 1, consistent with the smooth-solution error analysis [12]. The absolute differences at the production step *h* = 0.05 are below 1.6 *×* 10^−6^ in this test.

**Table 2:** Mesh convergence in the escape regime. The error is the largest absolute state difference at *τ* = 20 relative to the *h* = 0.00625 computation.

| $\alpha$ | $h$ | Maximum state error | Empirical order |
| --- | --- | --- | --- |
| 0.7 | 0.200 | $1.6892 \times 10^{-5}$ | — |
| | 0.100 | $5.2370 \times 10^{-6}$ | 1.690 |
| | 0.050 | $1.5954 \times 10^{-6}$ | 1.715 |
| | 0.025 | $4.6141 \times 10^{-7}$ | 1.790 |
| 1.0 | 0.200 | $2.0900 \times 10^{-5}$ | — |
| | 0.100 | $5.2168 \times 10^{-6}$ | 2.002 |
| | 0.050 | $1.2897 \times 10^{-6}$ | 2.016 |
| | 0.025 | $3.0718 \times 10^{-7}$ | 2.070 |

## 5 Results

### 5.1 Fractional order changes transients, not equilibria

Figure 1 shows untreated trajectories for *α* ∈ *{*0.5, 0.6, …, 1.0*}*. In the escape regime, the tumour moves toward the coexistence equilibrium. At *τ* = 100, *T* ranges from 0.2105 for *α* = 0.5 to 0.8909 for *α* = 1. This large finite-horizon difference is a difference in convergence speed: every curve has the same equilibrium *T* ^∗^ = 0.8914. It would therefore be incorrect to describe the smaller *T* (100) at low *α* as a new asymptotic tumour-control state.

**Figure 1:**
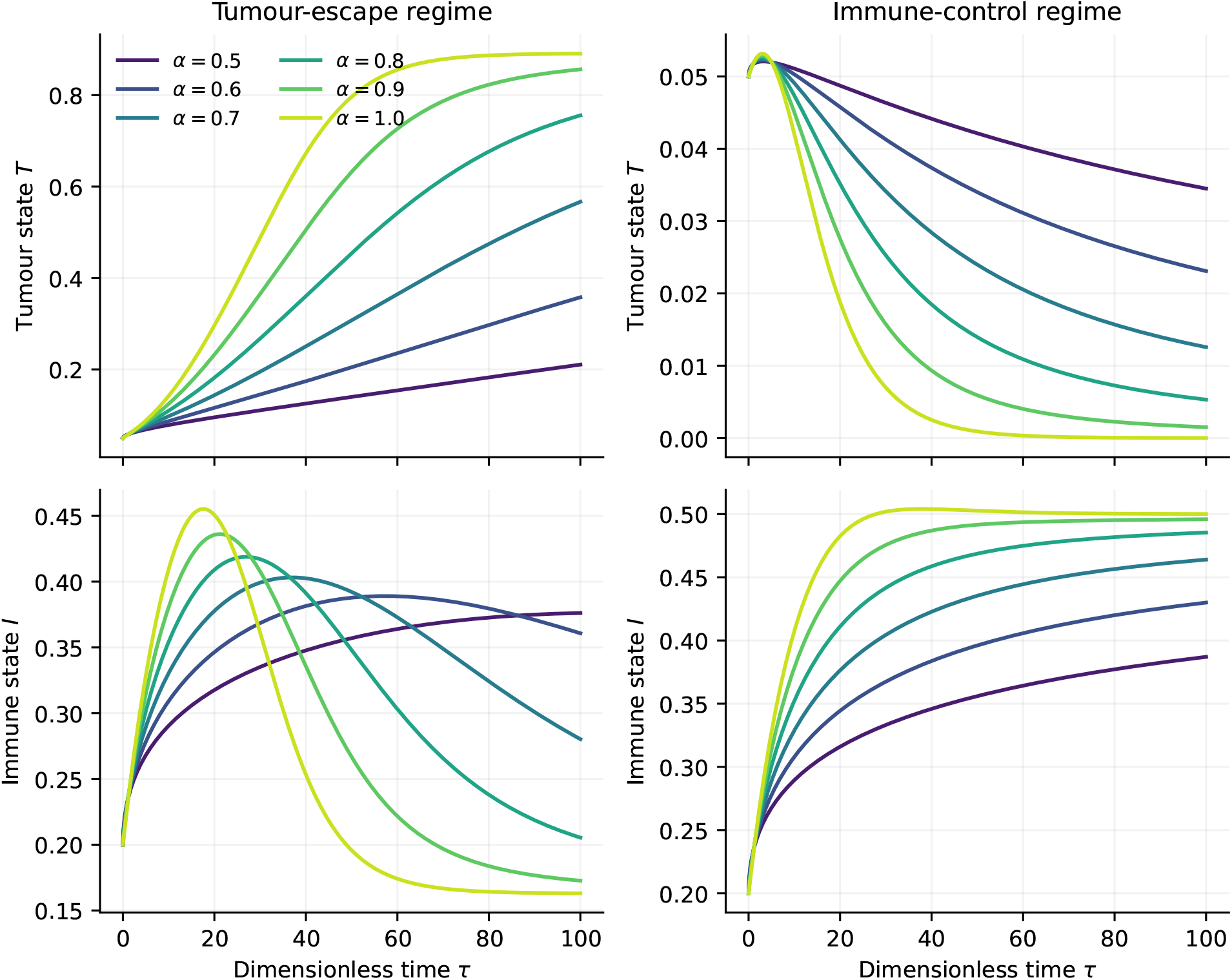
Untreated tumour and immune trajectories in the tumour-escape and immune-control regimes. The horizontal axis is dimensionless time. Lower fractional order slows approach to the corresponding equilibrium; it does not move that equilibrium.

In the immune-control regime, the tumour-free equilibrium is stable. Here the same time-scale effect appears in the opposite direction: *T* (100) = 0.03449 for *α* = 0.5, compared with 6 *×* 10^−6^ for *α* = 1. Smaller *α* preserves the transient tumour state for longer even though the asymptotic state is tumour-free in both cases. Table 3 summarizes the terminal values.

**Table 3:** Untreated terminal states at *τ* = 100.

| Regime | $\alpha$ | $T(100)$ | $I(100)$ | $N(100)$ |
| --- | --- | --- | --- | --- |
| Escape | 0.5 | 0.210515 | 0.376146 | 0.902496 |
|  | 0.7 | 0.566736 | 0.280254 | 0.659943 |
|  | 0.9 | 0.856428 | 0.172662 | 0.324899 |
|  | 1.0 | 0.890881 | 0.163060 | 0.200443 |
| Control | 0.5 | 0.034489 | 0.387044 | 0.978456 |
|  | 0.7 | 0.012564 | 0.463992 | 0.985448 |
|  | 0.9 | 0.001500 | 0.495932 | 0.997369 |
|  | 1.0 | 0.000006 | 0.500058 | 0.999941 |

### 5.2 Fractional bolus clearance

Figure 2 confirms the analytical result (8). Every concentration curve begins at and attains its maximum at *C* = 1 when *τ* = 0. Lower orders cross *C* = 0.5 slightly earlier in this parameterization, but decay much more slowly at late time. Accordingly, the finite-window area 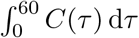 rises from 1.0006 at *α* = 1 to 7.8172 at *α* = 0.5 (Table 4). The area is truncated at *τ* = 60 because the infinite-horizon value does not exist for *α <* 1.

**Table 4:** Fractional clearance metrics. AUC is reported only on the finite interval [0, 60].

| $\alpha$ | $C_{\max}$ | Time of maximum | Half-time | AUC<sub>0:60</sub> |
| --- | --- | --- | --- | --- |
| 0.5 | 1.000 | 0 | 0.5938 | 7.8172 |
| 0.6 | 1.000 | 0 | 0.6072 | 5.4305 |
| 0.7 | 1.000 | 0 | 0.6243 | 3.6762 |
| 0.8 | 1.000 | 0 | 0.6442 | 2.4319 |
| 0.9 | 1.000 | 0 | 0.6676 | 1.5755 |
| 1.0 | 1.000 | 0 | 0.6936 | 1.0006 |

**Figure 2:**
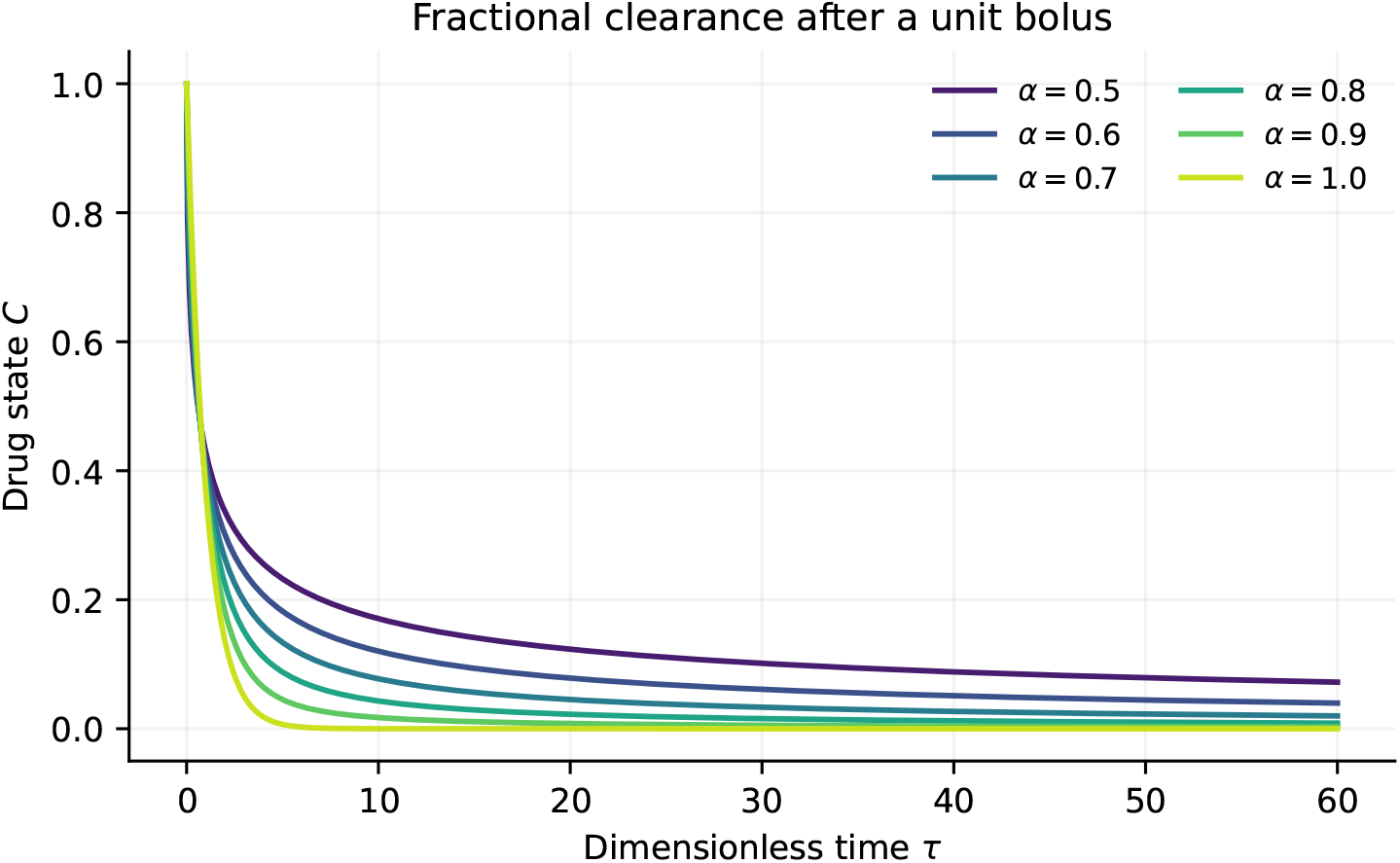
Drug state after a unit bolus, *C*(0) = 1, with *u* = 0. Smaller *α* produces a faster initial fall in this scaling but a much heavier late tail.

### 5.3 Coupled response to a unit bolus

Figure 3 shows the coupled escape-regime trajectories following *C*(0) = 1. Relative to the untreated trajectory at the same *α*, the tumour state at *τ* = 100 is reduced by 70.8% for *α* = 0.5, 57.4% for *α* = 0.7, 7.5% for *α* = 0.9, and 0.2% for *α* = 1. These percentages are mathematically reproducible but should not be called a therapeutic index. At *α <* 1, the drug state has the non-integrable tail in (9); the model therefore continues to apply weak drug pressure indefinitely. The late separation of the curves is a structural consequence of the clearance equation, not evidence that stronger fractional memory improves chemotherapy.

**Figure 3:**
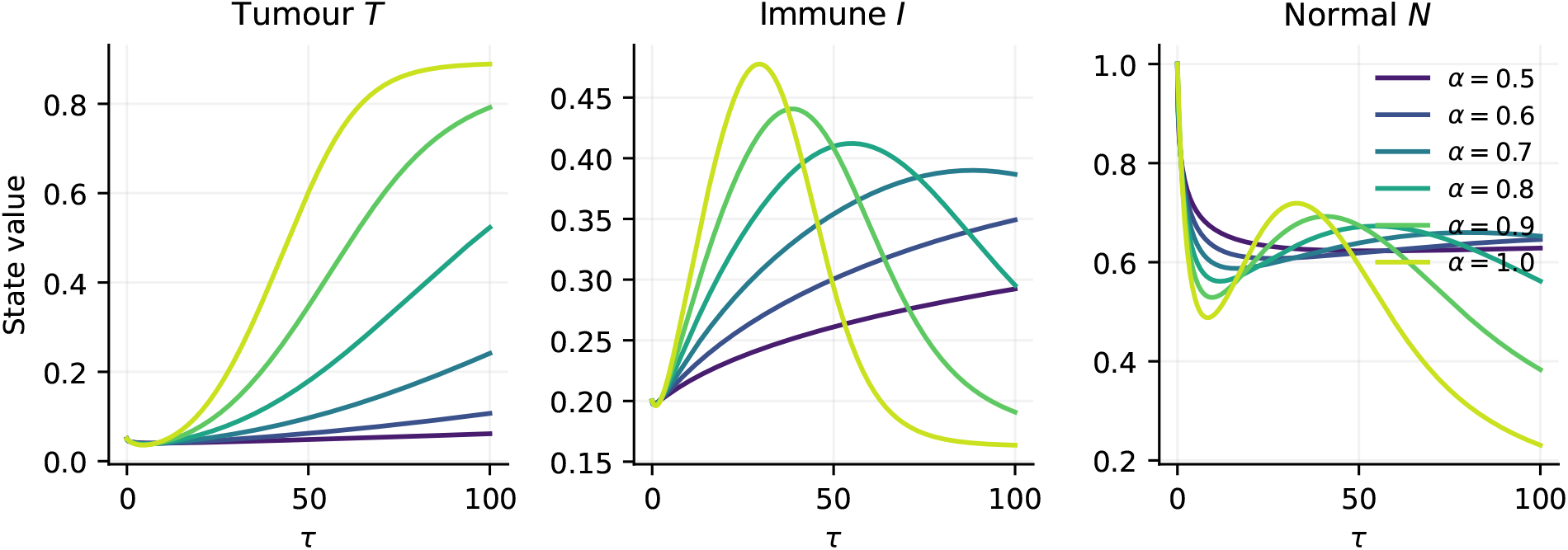
Coupled response to a unit bolus in the tumour-escape regime. Persistent suppression at low *α* is driven by the heavy drug tail and should not be interpreted clinically.

The model also predicts state-dependent toxicity. Minimum *N* values over 0 ≤ *τ* ≤ 100 are 0.6228, 0.5874, 0.3833, and 0.2308 for *α* = 0.5, 0.7, 0.9, and 1, respectively. These numbers cannot be mapped to clinical toxicity because the normal-cell state, time scale, and drug coefficient have not been calibrated.

## 6 Discussion

The calculations support a restrained interpretation of fractional-order tumour models. The Caputo order changes the temporal weighting of past rates and can substantially change finite-horizon trajectories. In the present commensurate system, however, it does not change equilibrium locations. A report of “final burden” at an arbitrary stopping time can therefore conflate slow fractional relaxation with a different asymptotic outcome. Both the escape and control regimes demonstrate this point: decreasing *α* preserves the initial condition longer, whether that means temporarily delaying tumour escape or temporarily delaying tumour elimination.

The pharmacokinetic equation requires even greater care. The exact solution proves that the concentration peak after a bolus occurs at the initial time. A delayed peak would require absorption, infusion, or another compartment. More importantly, the power-law tail gives an infinite-horizon AUC that diverges for *α <* 1. Heavy-tailed kinetics may be appropriate for some anomalous transport processes, and fractional pharmacokinetic models can be fitted successfully when their structure is chosen and validated carefully [7–9]. The simple equation used here is nevertheless unsuitable for unqualified claims about total exposure or therapeutic advantage.

Dimensionless formulation resolves, but does not eliminate, the scaling problem. Once a reference time *t*_ref_ is chosen, a physical derivative satisfies 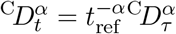. Changing *α* while keeping the same printed dimensional rate constants silently changes the model. A study intended to compare days or hours must specify how each coefficient varies with *α*, or estimate those coefficients separately for each order from data.

The present analysis also illustrates why statistical model-comparison claims require more than simulated trajectories. AIC, BIC, cross-validation, and confidence intervals need an identified dataset, an observation model, a fitted likelihood or loss, and a documented resampling procedure. None is available for this illustrative system. We therefore report deterministic numerical accuracy rather than fabricated inferential precision.

## 7 Limitations

This work is a theoretical and computational analysis, not a clinical study. The state variables and coefficients are dimensionless and are not calibrated to a specific cancer, drug, or patient population. A single common order *α* is imposed on all four equations, although distinct biological processes need not share one memory law. The model omits spatial heterogeneity, immune subtypes, resistance, stochasticity, drug absorption, distribution compartments, and discrete dosing events. Local stability does not by itself establish global convergence. Finally, the *O* (*n*^2^) history cost of the direct predictor–corrector implementation limits long-time fine-grid calculations; fast convolution methods would be preferable for large studies.

## 8 Conclusion

A dimensionless fractional tumour–immune–chemotherapy model can be analysed and simulated reproducibly, but its outputs require careful interpretation. The correct tumour-invasion number is *ℛ*_*T*_ = *r*_*T*_ *d*_*I*_*/*(*a*_*T*_ *s*_*I*_), with the tumour-free equilibrium stable when ℛ_*T*_ *<* 1. Equilibrium locations do not depend on the common Caputo order. Lowering *α* primarily slows relaxation and can make a finite-time state look better or worse depending on the underlying equilibrium. For the one-equation drug model, a unit bolus peaks at time zero and has an algebraic, non-integrable concentration tail when *α <* 1. Apparent late therapeutic gains in the coupled simulations are therefore model consequences, not clinical predictions. Future work should fit a dimensionally explicit model to longitudinal data and test whether a fractional memory law improves out-of-sample prediction over integer-order and distributed-delay alternatives.

## Code and data availability

The accompanying source package contains the Python script used to regenerate every figure and numerical table. No external, patient-level, or experimental data were used. Running `python code/reproduce.py` from the source directory recreates the contents of the `figures` and `results` directories.

## Ethics statement

Not applicable. This theoretical and computational study used no human participants, patient data, animals, or biological samples.

## Use of generative AI

Generative AI tools assisted with language editing and code drafting. The authors are responsible for verifying the mathematics, numerical results, references, and final submitted text.

## A Jacobian matrices

For *C* = 0, the Jacobian at a general state is

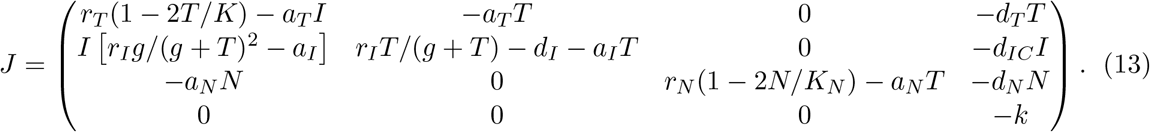

At *E*_0_, this matrix is lower block triangular and has the eigenvalues stated in Section 3.2. At a positive equilibrium, equations (6)–(7) can be used to simplify the diagonal entries before numerical evaluation.

## B Reproducibility checklist

The package records the software-independent model specification, all coefficient values, initial conditions, tested fractional orders, time horizon, and step size in results/run manifest.json. Machine-readable CSV files contain the values used in Tables 2–4. Vector PDF and high-resolution PNG versions of every figure are generated. The solver does not fall back to an integer-order approximation for *α <* 1, and no stochastic intervals are reported because no sampling distribution or observational dataset is part of the study.

